# Decorin inhibits glucose-induced lens epithelial cell apoptosis via suppressing p22^phox^-p38 MAPK signaling pathway

**DOI:** 10.1101/800680

**Authors:** Shanshan Du, Jingzhi Shao, Dandan Xie, Fengyan Zhang

**Author notes:** contributed equally to this study and should be considered as co-first authors (Shanshan Du, Jingzhi Shao). Corresponding author. Email address (Fengyan Zhang).

## Abstract

**Purpose:** To determine the effect of decorin on oxidative stress and apoptosis of human lens epithelial (HLE) cells under high glucose condition.

**Methods:** HLE cell line (HLEB3) was incubated in normal glucose (5.5 mM) or high glucose (60 mM) medium. Decorin (50 nM) was applied 2 hours before high glucose medium was added. Apoptosis detection was executed by flow cytometry and western blotting (analysis of bcl-2 and bax). Oxidative stress level was measured by the generation of reactive oxygen species (ROS), glutathione peroxidase (GSH) and superoxide dismutase (SOD). P38 mitogen-activated protein kinase (MAPK) phosphorylation, the expression of p22^phox^ of HLE cells and human lens anterior capsules were detected by western blotting. Small interfering RNA transfection to p22^phox^ and p38 MAPK was also carried out on HLEB3.

**Results:** High glucose caused HLE cells oxidative stress and apoptosis exhibiting the increase of apoptotic cells and ROS production and decrease of bcl-2/bax ratio, GSH/GSSG ration and SOD activity. P22^phox^ and phospho-p38 MAPK were upregulated in high glucose treated HLEB3 cells. Knocking down p22^phox^ or p38 by siRNAs can reduce high glucose induced cell apoptosis and oxidative stress level. Silencing p22^phox^ by siRNA can downregulate p38 MAPK activation. Decorin can inhibit the apoptosis, oxidative stress level and the induction of p22^phox^ and p-p38 of HLEB3 induced by high glucose. Furthermore, the expression of p22^phox^ and p38 were found significantly increased in lens anterior capsules of diabetic cataract patients compared to that of normal age-related cataract patients.

**Conclusions:** Results showed that p22^phox^-p38 pathway may be particepated in high glucose induced lens epithelial cell injury, decorin may inhibit the high glucose induced apoptosis and oxidative stress injury by suppressing this pathway in part.

## Introduction

Diabetic cataract is one of the most important complications of diabetes (1). Oxidative stress induced by high glucose plays a pivotal role in the mechanism of diabetic cataract. Oxidation and aggregation of protein in lens epithelium cells, which led to lens opacity, are caused by high glucose (2). Apoptosis and oxidative stress, which participated in the formation of diabetic cataract, occurred when human lens epithelial (HLE) cells exposing for 24 h to the dondition of high glucose (3).

Decorin, which is a small leucine-rich proteoglycan, has been found to negatively regulate a variety of cellular functions when binding to extracellular matrix components or cell surface receptors (4, 5). Overexpression of decorin could restrain angiogenesis mediated by tumor cell by suppresssing the production of vascular endothelial growth factor (VEGF) (6). Administration of decorin into corneal stroma could inhibit neovascularization of cornea in rabbit model (7). In the post-traumatic brain injuries (TBI) rat cerebrum, it is reported that decorin increased the activity of superoxide dismutase (SOD), glutathione peroxidase (GSH) and prevented oxidative stress injury and apoptosis (8, 9). Preliminary data of our team showed that decorin can inhibit retina pigmentosa epithelia (RPE) barrier disruption under diabetic condition through suppression of p38 mitogen-activated protein kinase (MAPK) activation (10). However, the influence of decorin on diabetic cataract has not been studied yet.

In the current study, the influence and potential mechanism of decorin was excuted on high glucose-induced oxidative stress and apoptosis in HLE cells. Apoptosis, reactive oxygen species (ROS), SOD and GSH were determined. Besides, p38 MAPK phosphorylation and the expressed level of p22^phox^ of HLEB3 and human lens anterior capsules were evaluated. Furthermore, the role of p22^phox^ and p38 MAPK were evaluated in mediating the oxidative stress caused by high glucose.

## Materials and Methods

### 2.1. Antibodies and chemical agents

Mouse anti-bcl-2 (Abcam, Cambridge, UK), mouse anti-bax (Abcam, Cambridge, UK), mouse anti-β-actin (Proteintech, Chicago, Illinois, USA), rabbit anti-p38 (Abcam, Cambridge, UK), rabbit anti-phospho-p38 (Thr180/Tyr182, Abcam, Cambridge, UK), mouse anti-p22phox (Gentex, Irvine, USA), goat anti-mouse (Proteintech, Chicago, Illinois, USA) and goat anti-rabbit (Proteintech, Chicago, Illinois, USA) antibodies, recombinant human decorin (R&D Systems, Minneapolis, MN, USA), apoptosis detection kit (Beyotime Inst. Biotech, Haimen, China), Cell Counting Kit-8 (CCK-8, Dojindo, Japan).

### 2.2 Human lens anterior capsules

The human lens anterior capsular tissues were obtained from the eye-tissue bank. All procedures for collecting the anterior capsules followed the guidelines for the use of human materials. The protocols used in the paper were approved by the Ethics Committee of the First Affiliated Hospital of Zhengzhou University. All patients offered written reports of informed consent. Twenty-six cataract patients (thirty eyes) with type 2 diabetes mellitus were included as the diabetic cataract group, and twenty-three cataract patients (thirty eyes) without any systemic or ocular disease were included as the senile cataract group (control group) (Table 1).

**Table 1.** Information of cataract patients

|  | Diabetic Cataract Patients |  | Senile Cataract Patients |  |
| --- | --- | --- | --- | --- |
| | Mean ± SD | Range | Mean ± SD | Range |
| Sample size (M:F) | 26 (11:15) |  | 23 (11:12) |  |
| Eye number (R:L) | 30 (13:17) |  | 30 (14:16) |  |
| Diabetes, year | 9.58 ± 6.99 | 0.25 - 20 | | |
| Age, yr | 65.93 ± 7.84 | 46 - 89 | 74.43 ± 8.66 | 58 - 91 |
| Cataract grade | 2.93 ± 0.68 | 1 - 4 | 2.79 ± 0.86 | 1 - 4 |
| Hemoglobin A1c, % | 8.02 ± 0.68 | 7 - 10 | | |
| Receive treatment | 100% |  |  |  |

### 2.3 Human lens epithelial cell cultures

Human lens epithelial cell line (HLEB3) was a gift of Professor Hu in Henan Unicersity. This cell line was maintained in Dulbecco’s modified Eagle’s medium (DMEM with 5.5mM D-glucose) (Sigma, USA) containing 10% fetal bovine serum (FBS) (Genial, USA) and 1% antibiotics (penicillin G 110 U/ml, streptomycin 100μg/ml, solarbio, China) at 37 °C in the presence of 5% CO_2_. The cells were passaged every three days. Cells for 2 to 5 passages were used.

HLEB3 cells were seeded in 6-well plates with normal culture. When cell density reached about 50%, the cells were exposed to serum starvation (0.5% FBS) for 12 hours. Then the cells were divided into four groups: control group, high glucose group, decorin group and mannitol group. Cells in high glucose group were exposed to glucose (60 mM, Sigma, USA) for 72 hours. Cells in decorin group were exposed to recombinant human decorin (50 nM) for 2 hours (11) and then treated with 60 mM glucose for 72 hours. Mannitol group (5.5 mM D-glucose plus 54.5mM mannitol) was designed to eliminate potential deviations by osmosis.

### 2.4 Cell Viability Assay

CCK-8 was used to measure cell viability. HLEB3 cells were seeded in 96-well plates at a density of 1×10^4^ cells per well for overnight. Then the medium was changed with medium (without FBS) containing 0, 50, 100 and 200nM of decorin. Plates were incubated for 24 h, then 10μl WST-8 [2-(2-methoxy-4-nitrophenyl)-3-(4-nitrophenyl)-5-(2,4-disulfophenyl)-2H-tetrazolium, monosodium salt] solution was added to each well. After incubating for 4 hours at 37°C, the plate was examined at 450nm using an iMark™ microplate absorbance reader (Bio-Rad, Hercules, USA). Number of living cells in each well was expressed as the value relative to the control.

### 2.5 Small Interfering RNA Transfection

Small Interfering RNA (siRNA) targeted to p22^phox^ or p38 MAPK was purchased from San Cruz Biotechnology (Dallas, Texas, USA). The siRNA or scramble control siRNA was transfected to HLEB3 cells following the kit’s protocol. After incubating for 6 hours with 100 nM siRNA, the medium was changed with high glucose as described above.

### 2.6 Flow Cytometry

Cell apoptosis was measured by flow cytometry with Annexin-V-FITC apoptosis detection kit. HLEB3 cells were collected and stained with Annexin-V-FITC for 10 min at room temperature. The cells were centrifuged and washed in PBS buffer. Then, the cells were resuspended in PBS containing propidium iodide (PI) and subjected to flow cytometer (BD^TMc^LSR II). Data were analyzed using Win MDI 2.9 software.

### 2.7 Measurement of intracellular ROS

IIntracellular ROS was tested by the densitometry measuring of 2’, 7’-dichlorofluorescein diacetate (DCFH-DA, Beyotime Inst. Biotech, Haimen, China) fluoresce. Under oxidative stress condition, DCFH-DA was oxidized and converted to fluorescent dichlorofluorescein (DCF), which could be measured by flow cytometer. Briefly, HLEB3 cells were incubated in diluted DCFH-DA for 20 minutes and then washed with serum-free DMEM. Then the cells was captured with fluorescence microscope or collected for measuring with BD Accuri C6 Flow Cytometer (Bioscience; USA) with excitation and emission wavelengths at 488 nm and 525 nm.

### 2.8 Detection of SOD and GSH

After harvested, cells were washed twice with PBS. Intracellular SOD activity and GSH/GSSG ratio were measured following the protocols of total SOD assay kit and GSH/GSSG assay kit (Beyotime Inst. Biotech, Shanghai, China). Then the plates were measured at 450nm and 412nm separately using an iMark™ microplate absorbance reader (Bio-Rad, Hercules, USA).

### 2.9 Western blotting

Western blotting analysis was performed as previously described [14]. Briefly, the equal amount of protein was separated in SDS-PAGE gel and transferred to polyvinylidene fluoride (PVDF) membrane. After incubating for 1 hour with 5% milk in TBST buffer (25 mM Tris-HCl, pH 7.4, 150 mM NaCl, 1% triton X-20) at room temperature, membranes were incubated with primary antibodies overnight at 4°C. Wshed in TBST buffer for three times, the membranes were incubated with horseradish peroxidase-conjugated secondary antibody. The membranes were incubated with electrochemiluminescence (ECL) buffer and signals were exposed to X-ray film.

For the examination of human lens anterior capsules, the anterior capsules were collected from patients who underwent cataract surgery. In order to get enough protein for analysis, ten anterior capsules from each group were mixed together and subjected to immunoblotting.

### 2.10 Statistical Analysis

Data from at least three independent experiments were represented as means ± standard error of the mean (SEM). Statistical analysis was carried out using SPSS 18.0 software. One-way ANOVA was used for comparisons among more than two groups. Student-Newman-Keuls test was used for comparisons between two groups. *P* value of < 0.05 was statistically significant.

## Results

### 3.1 Cell Viability Assay

Cell viability of HLEB3 cells in different concentration of decorin (0nM, 50nM, 100nM and 200nM) showed no significant difference when examined by CCK-8 assay (Figure. 1). Moreover, morphological appearance of HLEB3 cells was observed by phase contrast microscopy. We did not find obvious morphological changes at different concentrations of decorin (Figures not showed).

**Figure 1.**
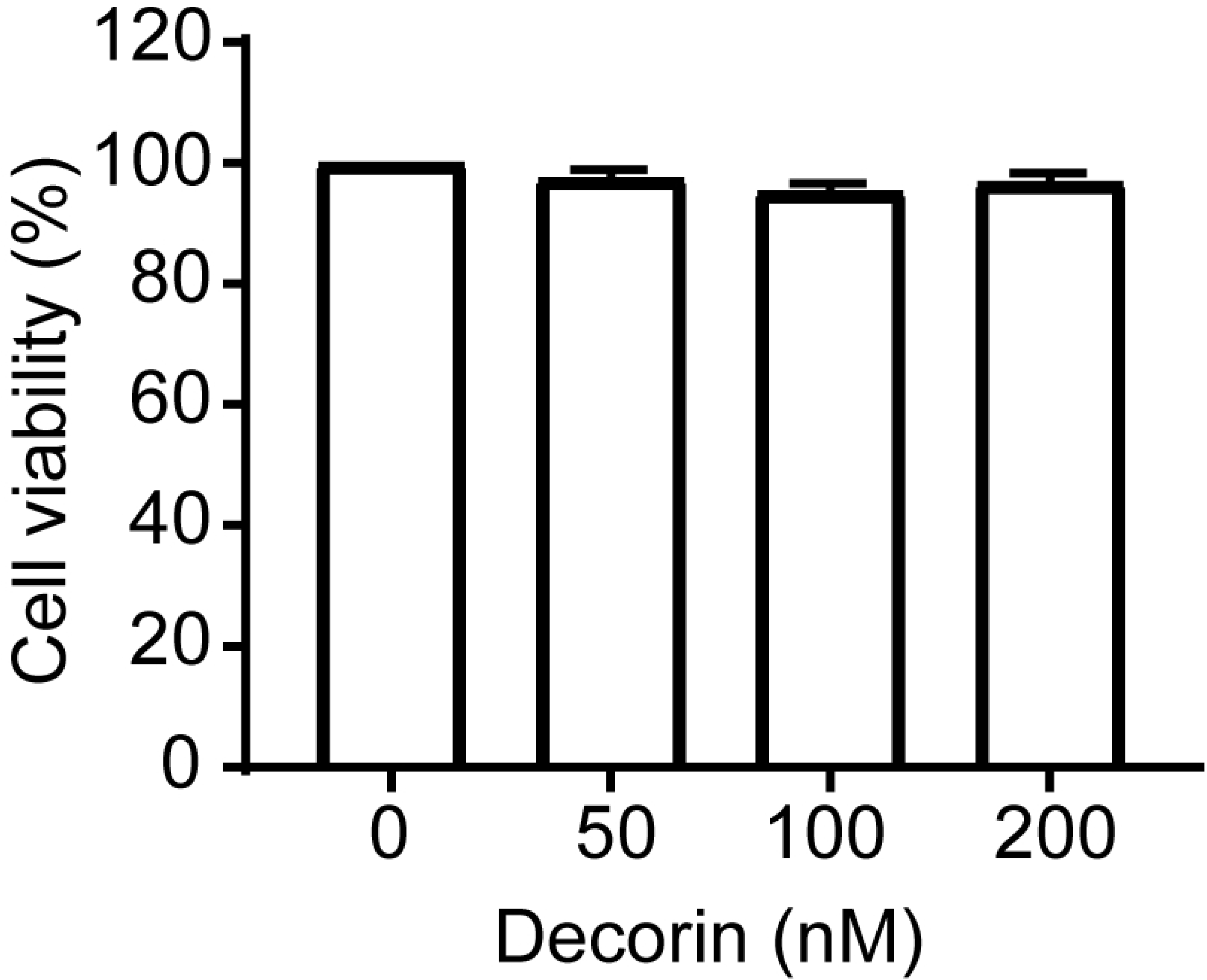
Cell viability of HLEB3 cells at different concentration of decorin as tested by CCK-8 assay. No significant differences in cell viability were found among the different concentration of decorin tested (*P* > 0.05). Bars are means ± SEM.

### 3.2 Decorin inhibits apoptosis of HLEB3 cells induced by high glucose

The influence of decorin on the apoptosis of HLEB3 cells are shown in Figure 2. The cells were distinguished into four groups (Figure 2A): unlabeled cells (viable cells), cells that stained with annexin V-FITC only (early apoptotic), cell that stained with PI only (necrotic), and cells that stained with both annexin V and PI (late apoptotic). The sum of late apoptotic cells and early apoptotic cells was defined as apoptotic cells. For the control group, the percent of apoptotic cells was 4.5% ± 0.6%. After treatment with high glucose, the apoptotic percentage increased to 9.9% ± 0.4% (*P* <0.05 vs the control). When cells were treated with 50 nM decorin before high glucose administration, the apoptotic percentage decreased to almost normal level (Figure 2A and 2B). Additionally, the western bolt results showed that high glucosemake the protein expression level of bax and increased and the protein expression level of bcl-2 of HLEB3 cells decreased. The densitometry quantitation indicated that the ratio of bcl-2 to bax decreased significantly in high glucose media compared to that of control. Decorin can reverse the expression ratio of bcl-2 to bax in high glucose-treated cells (Figure 2C and 2D).

**Figure 2.**
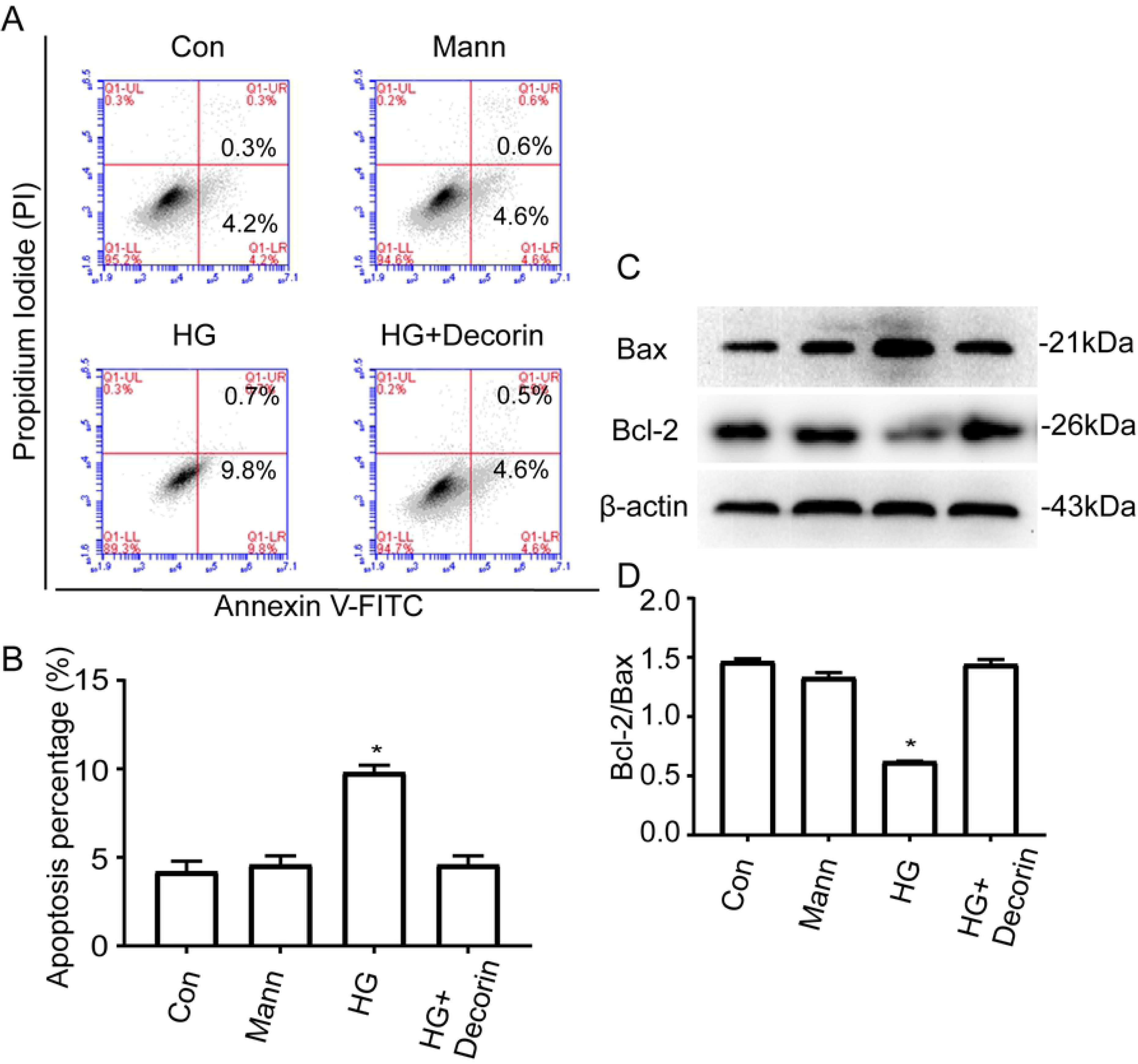
Effects of decorin on cell apoptosis. A: Quantitative analysis of HLEB3 cell apoptosis tested by flow cytometry with Annexin-V-FITC apoptosis detection kit. The upper right quadrants represented the late poptotic cells, and the lower right quadrants represented the early apoptotic cells. B: The sum of late apoptotic cells and early apoptotic cells was defined as apoptotic cells. The apoptotic cell percentage was statistically analyzed. C: Western blot analysis of bcl-2 and bax expression in HLEB3 cells. D: Optical density ratio of bcl-2 to bax. Decorin reduced high glucose-induced apoptosis of HLEB3 cells. Con, Control; Mann, Mannital; HG, high glucose. Data were expressed as mean ± SEM. \**P* < 0.05 vs the control group.

### 3.3 Decorin inhibits the oxidative stress response of HLEB3 cells under high glucose condition

Results indicated that ROS production was significantly increased in high glucose condition (Figure 3A and 3B), and the ratio of GSH/GSSG (Figure 3C) and the SOD activity of HLEB3 cells were decreased significantly (Figure 3D). Decorin can reverse this phenomenon induced by high glucose (Figure 3).

**Figure 3.**
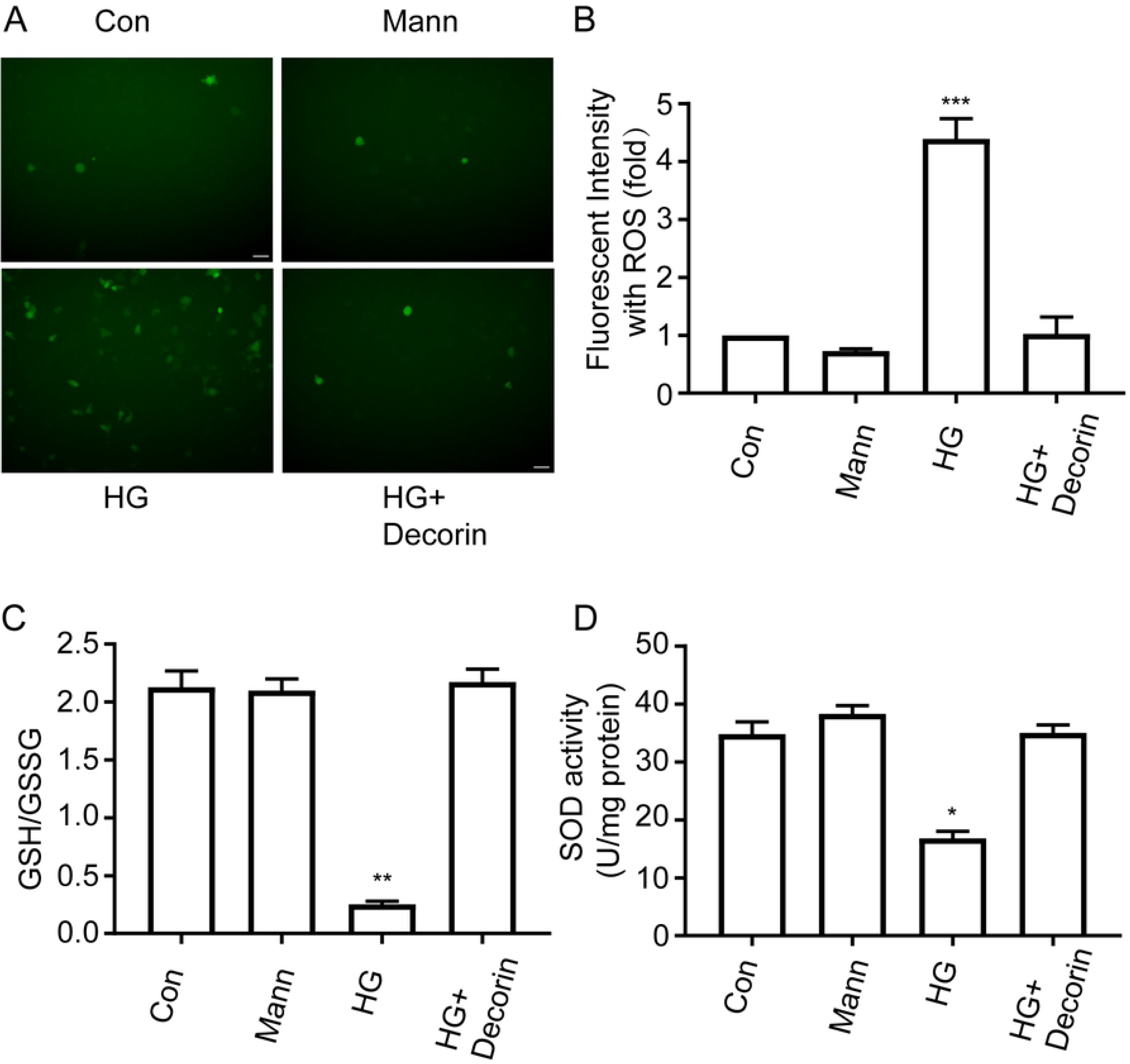
Alteration of intracellular oxidative stress. A-B: Fluorescent intensity with ROS level examined by DCFH-DA staining obvioused by fluorescence microscopy. Scale bars = 20 μm. C: GSH/GSSG ratio in the indicated groups. D: The activity of SOD in the indicated groups. Decorin reduced oxidative stress response of HLEB3 cells induced by high glucose. Con, Control; Mann, Mannital; HG, high glucose. Data were means ± SEM. \**P* < 0.05, \*\**P* < 0.01, \*\*\**P* < 0.001, compared to the control group respectively.

### 3.4 Decorin inhibits the activation of p38 and p22^phox^ in HLEB3 cells induced by high glucose

Western blot analysis was used to examine the activation of p38 and p22^phox^, which may participate in the effects of decorin on HLEB3 cells. Density analysis indicated that high glucose significantly increased the activation of p38 MAPK (Figure 4A and 4B) and the expression level of p22^phox^ protein (Figure 4C and 4D). When treated with 50 nM decorin, HLEB3 cells expressed decreased phosphorylation of p38 MAPK protein and the expression of p22^phox^ protein compared with high glucose group (Figure 4).

**Figure 4.**
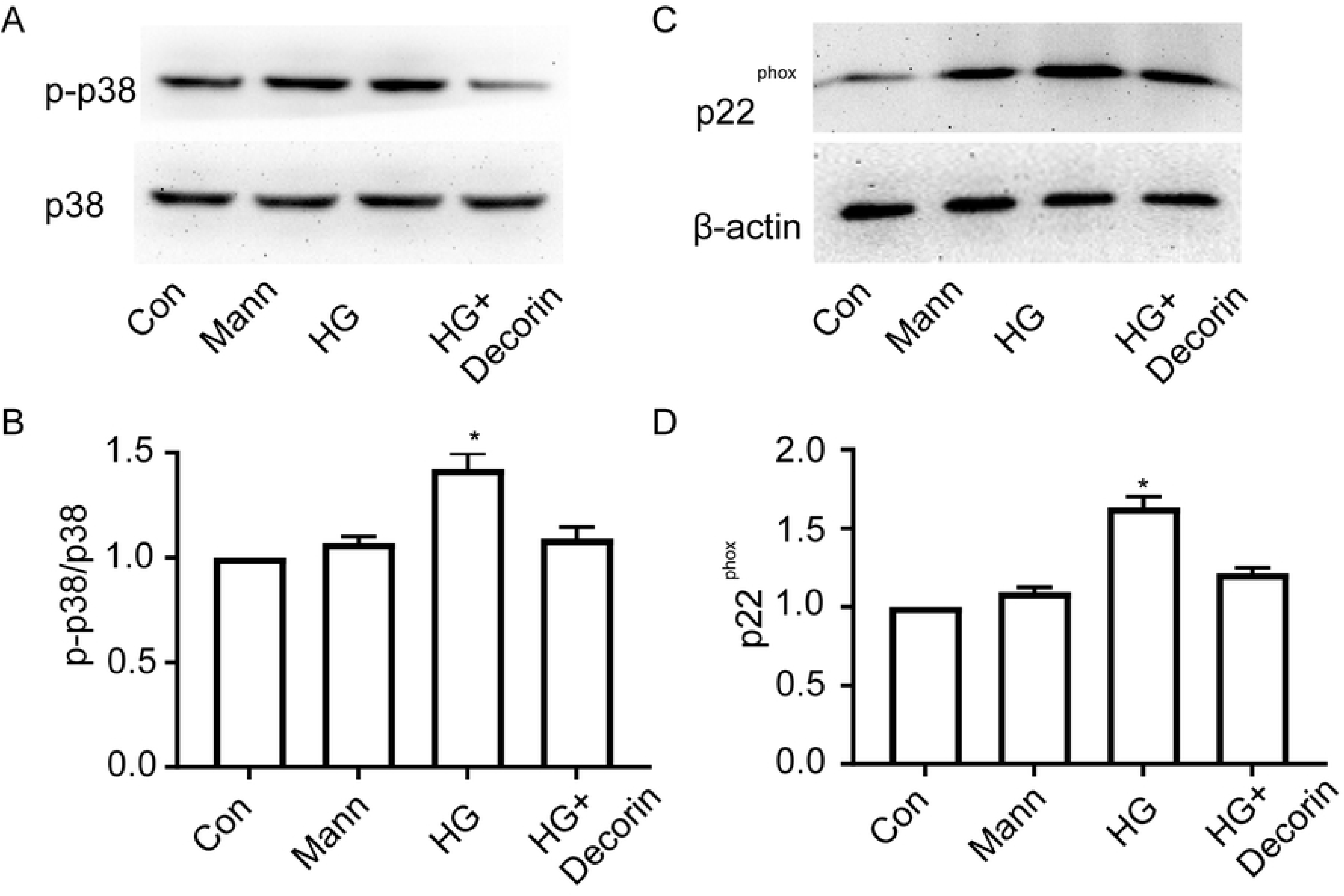
Effects of decorin on high glucose-induced activation of p38 and p22^phox^ in HLEB3 cells. A-B: p38 MAPK phosphorylation was presented by western blot. C-D: Western blot results of p22^phox^ expression. β-actin was regarded as an internal control. Decorin reduced the activation of p38 and the expression p22^phox^ in HLEB3 cells under high glucose condition. Con, Control; Mann, Mannital; HG, high glucose. Data were means ± SEM. \**P* <0.05 vs control group.

### 3.5 P38 MAPK is involved in regulating the oxidative stress and apoptosis of HLEB3 cells induced by high glucose

We do not know whether p38 MAPK was related to the high glucose-induced cell apoptosis and oxidative stress. To definited this, siRNA targeting p38 MAPK was used to transfected with HLEB3 cells. Western blot was used to examine the expression level of p38 MAPK. Results showed that the expression of p38 MAPK was significantly reduced by p38 siRNA (Figure 5A and Figure 5B). When cells were transferred with siRNA, flow cytometry analysis showed that the apoptotic percentage of p38 MAPK siRNA-transfected group was greatly reduced compared with control si RNA transfected group (*P* <0.05 vs the control) (Fig 5C and 5D). High glucose-induced expression level of bax and bcl-2 was reversed by p38 siRNA transfection (Figure 5E and Figure 5F). Furthermore, transfection of p38 siRNA significantly reduced the ROS production, increased the ratio of GSH/GSSG and increased the SOD activity in HLEB3 cells treated with high glucose (Figure 6).

**Figure 5.**
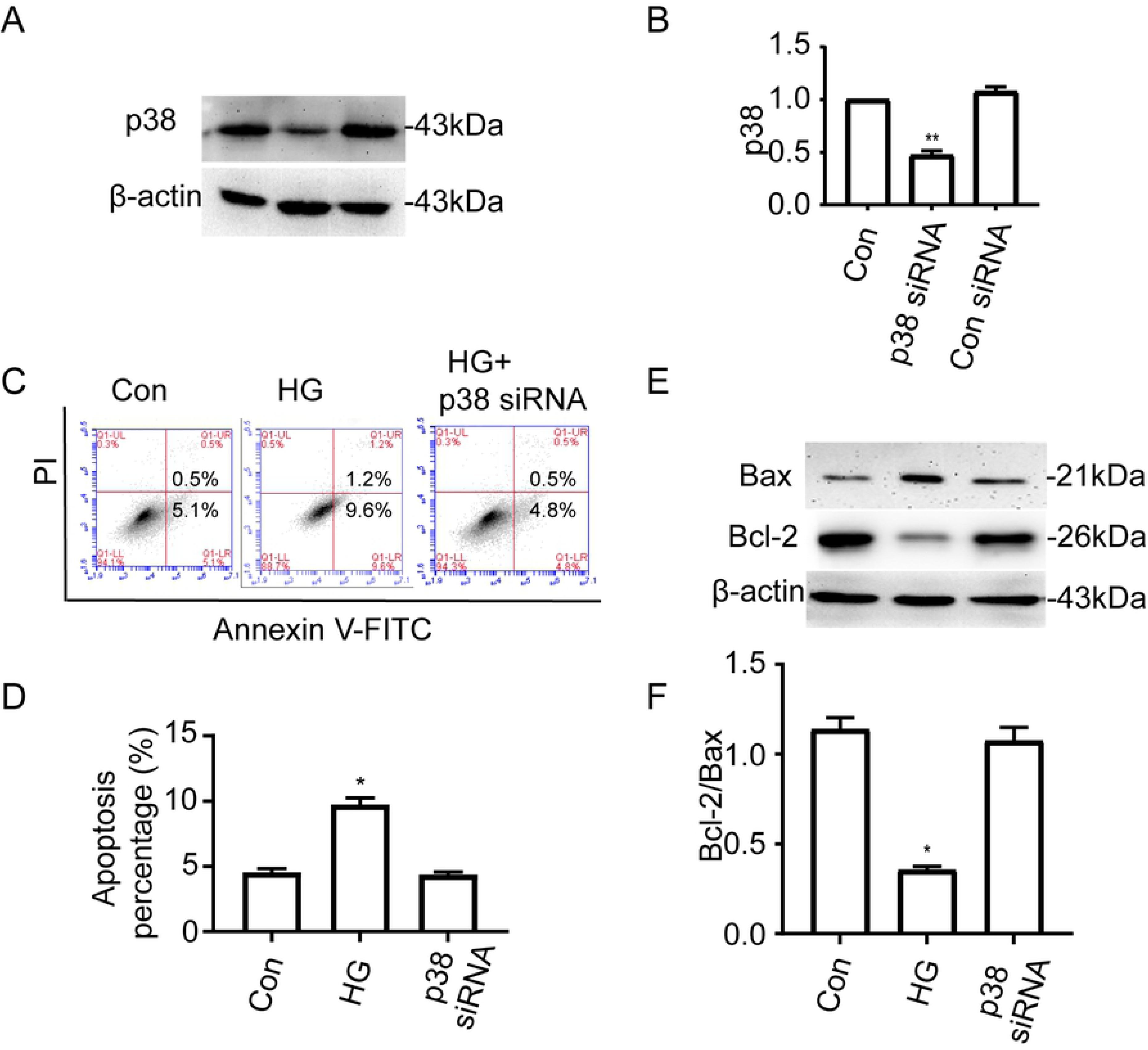
siRNA targeting p38 MAPK inhibited the apoptosis of HLEB3 cells induced by high glucose. A-B: Cells transfected with p38 MAPK siRNA resulted in a significant reduction in expression level of p38 protein. Cells transfection with control siRNA did not affect the expression level of p38 protein. C. Quantitative analysis of HLEB3 cell apoptosis was tested by flow cytometry with an Annexin-V-FITC apoptosis detection kit. The upper right quadrants represent the late apoptotic cells. The lower right quadrants represent the early apoptotic cells. D. The sum of late apoptotic cells and early apoptotic cells was defined as apoptotic cells. The apoptotic percentage was statistically analyzed. E: Western blot analysis of bcl-2 and bax expression. F: Optical density ratio of bcl-2 to bax. Con, Control; HG, high glucose. Data were expressed as means ± SEM. \**P* <0.05 vs control group, \*\**P* <0.01 vs control group.

**Figure 6.**
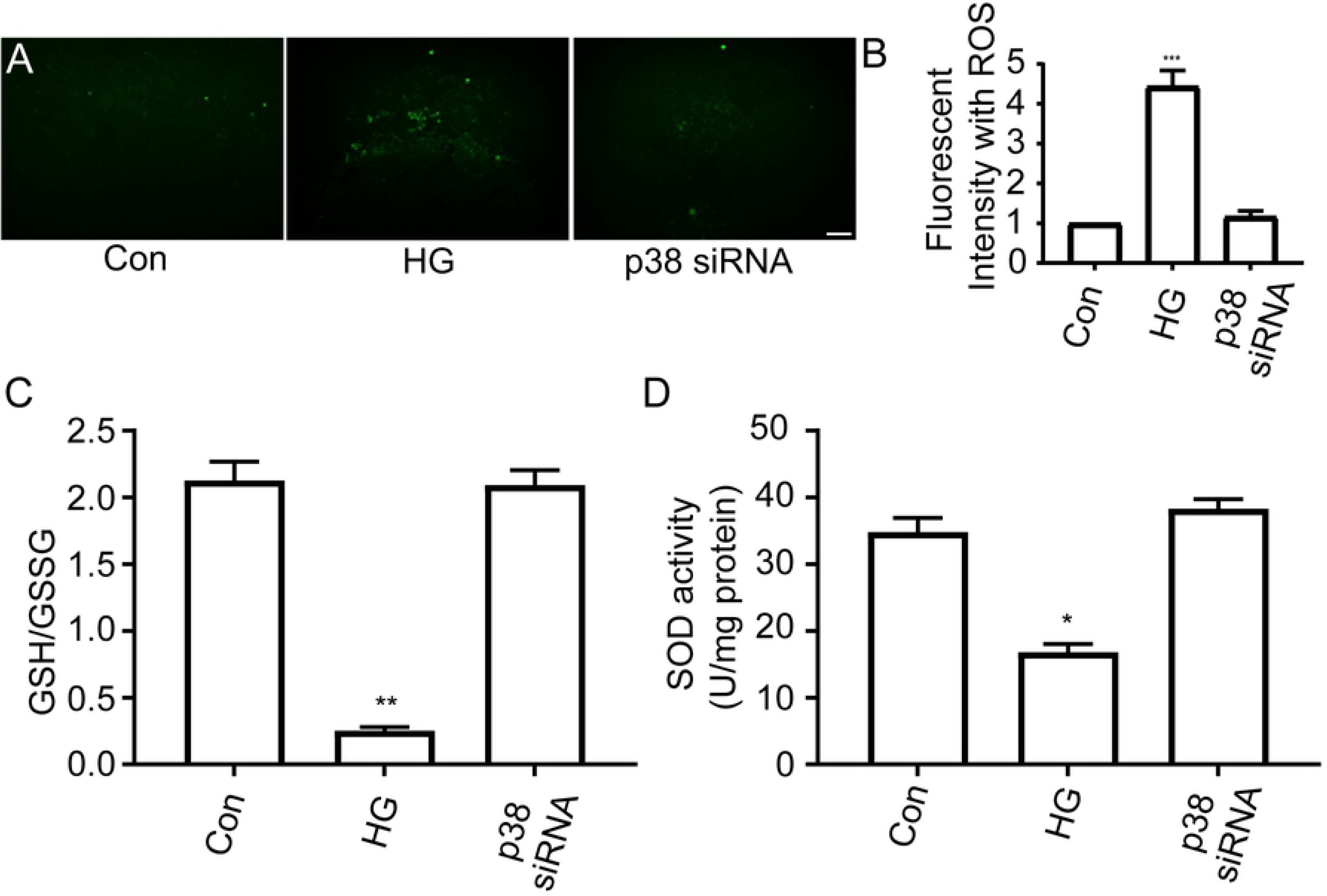
Alteration of intracellular oxidative stress after p38 MAPK siRNA transfection. A-B: Fluorescent intensity with ROS level was examined by DCFH-DA staining with fluorescence microscopy. Scale bars = 20 μm. C: GSH/GSSG ratio in the indicated groups. D: The activity of SOD in the indicated groups. High glucose-induced increase of oxidative stress was nearly inhibited in cells transfected with p38 MAPK siRNA. Con, Control; HG, high glucose. Data were means ± SEM. \**P* < 0.05, \*\**P* < 0.01, \*\*\**P* < 0.001, compared to the control group respectively.

### 3.6 p22^phox^-p38 MAPK pathway may mediate apoptosis and oxidative stress of HLEB3 cells induced by high glucose

To determine whether there is any possible association between p38 MAPK and p22^phox^ in the regulation of oxidative stress induced by high glucose, siRNA targeting p22^phox^ was transfected with HELB3 cells. Western blot was used to examine the expression level of protein. Results indicated that the expression of p22^phox^ was significantly reduced by p22^phox^ siRNA (Figure 7A and Figure 7B). Transfection of p22^phox^ siRNA can decrease the expression of phospho-p38 in cells that under high glucose condition (Figure 7C and Figure 7D). These results suggest that p22^phox^ regulates the activation of p38 MAPK and may mediate high glucose-induced apoptosis and oxidative stress of HLEB3 cells.

**Figure 7.**
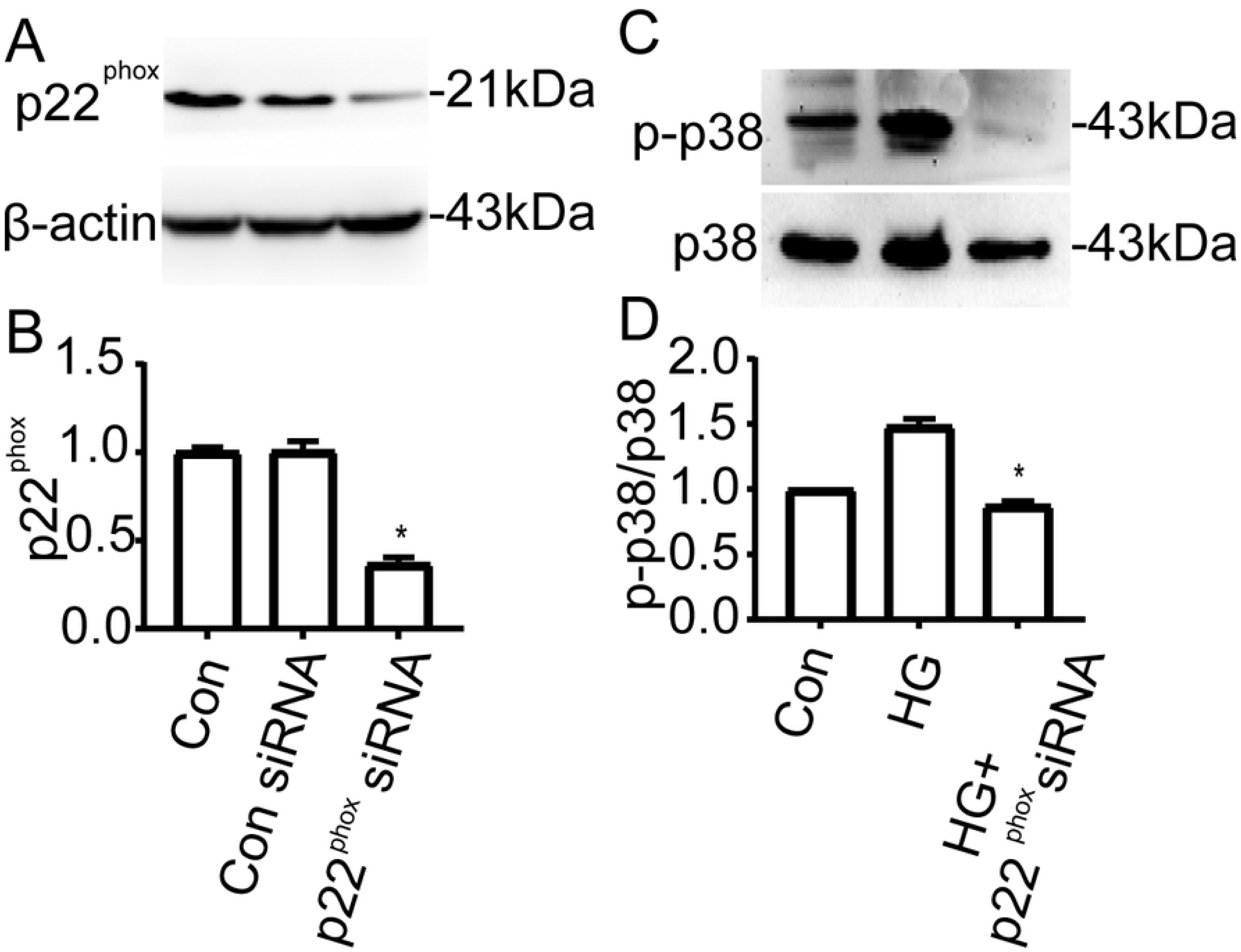
Possible association between p38 MAPK and p22^phox^ in the regulation of oxidative stress induced by high glucose. A-B: Transfection with p22^phox^ siRNA resulted in a significant reduction in p22^phox^ protein expression. Transfection with control siRNA did not affect p22^phox^ expression. C-D: Transfection with p22^phox^ siRNA resulted in obvious reduction in phorsphor-p38 MAPK expression. P22^phox^ regulates the activation of p38 MAPK. Con, Control; HG, high glucose. Bars were means ± SEM. β-actin was used as an internal control. \**P* < 0.05 vs control group.

### 3.7 The increased expression of p22^phox^ and phospho-p38 in diabetic lens anterior capsules

Additionally, we examined the expression of phosphor-p38 MAPK and p22^phox^ in diabetic lens anterior capsules. Results indicated that the expression level of phospho-p38 was significantly increased in anterior capsules from diabetic cataract patients vursus to that from simple senile cataract patients (Figure 8A and Figure 8B). Similar results were also observed for the expression of p22^phox^ (Figure 8C and Figure8D). These results suggest that p22^phox^-p38 pathway may participate in modulating the oxidative stress in diabetic lens.

**Figure 8.**
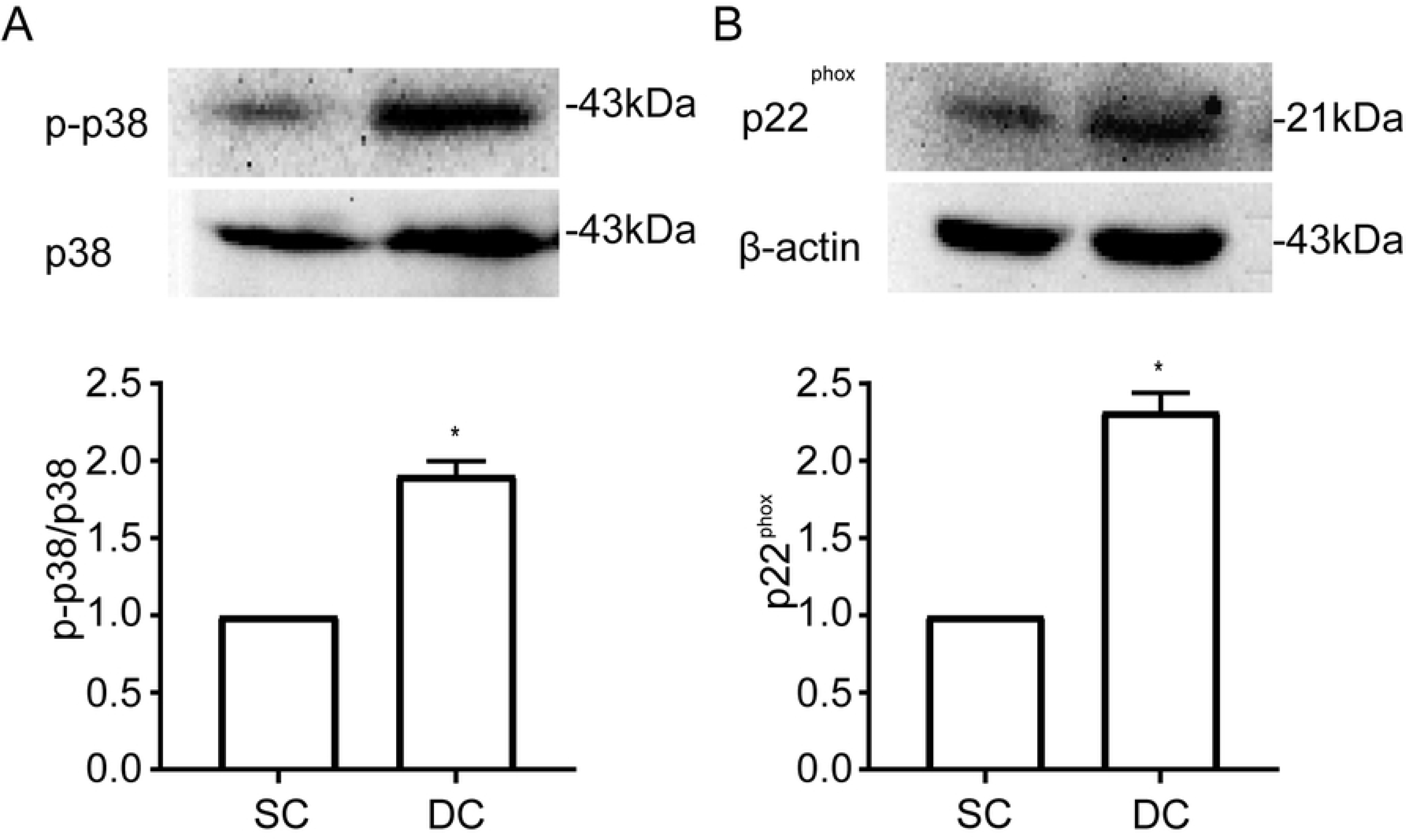
Expression of p22^phox^ and phospho-p38 in human lens anterior capsules. A-B: Western blot analysis of p38 MAPK expression. C-D: Western blot analysis of p22^phox^ expression. The expression of phospho-p38 and p22^phox^ were both significantly increased in anterior capsules of diabetic cataract patients vursus to that of simple senile cataract patients. SC, senile cataract; DC, diabetic cataract. β-actin was regarded as an internal control. Bars were means ± SEM. \**P* < 0.05 vs SC group.

## Discussion

High glucose-induced oxidative stress is an important cause of diabetic cataract (2). The human lens epithelial cell line (HLEB3) were widely used to do research (12–14). Here, using the lens epithelial cell line, we found that high glucose can directly induce the oxidative stress and cell apoptosis. P38 and p22phox are important signaling factors that regulate the high glucose-induced oxidative stress and apoptosis, and p22phox works upstream of p38 MAPK. The administration of decorin can inhibit the cellular oxidative stress and cell apoptosis induced by high glucose. Decorin can also inhibit high glucose-induced p22phox expression and p38 MAPK activation. These results implied that decorin exhibits the potential therapeutic role to diabetic cataract.

P22phox is a subunit of nicotinamide adenine dinucleotide phosphate (NAPDH) oxidase (NOX) that is responsible for ROS production in mitochondria at oxidative stress condition(15). p22phox is upregulated in diabetic endothelial cells and enhance the injury of endothelial cells by high concentration of glucose(16). The increase of p22phox expression induces the oxidative stress injury via activation of p38 MAPK(17). Our data showed that this signaling pathway also worked in diabetic lens epithelial cell injury. The supporting data are: 1) high glucose could directly induce ROS production, downregulate the ratio of GSH/GSSG and inhibit the SOD activity that lead to increase of the lens epithelial cell apoptosis; 2) high glucose can induce the expression of p22phox and p38 MAPK activation. Knocking down p22phox downregulates the high glucose-induced p38 MAPK activation, and knocking down p38 MAPK can inhibit high glucose-induced ROS production and cell apoptosis. The viewpoint that ROS is a key inducer of cell apoptosis or necrosis in divergent tissues has been widely accepted (15); 3) additionally, we examined that the human lens anterior capsules from diabetic cataract patients had higher expression of phospho-p38 and p22phox compared with that from senile cataract patients. These results suggested that p38 MAPK is downstream of p22phox in the regulation of ROS production in lens epithelial cells at diabetic stress condition.

Decorin is a small leucine-rich proteoglycan constituting the extracellular matrix binding to collagen type I fibrils(4, 5). In addition, decorin is a signaling molecule involving in regulating the angiogenesis, anti-apoptosis, tumor growth and inflammation(6–9). Our data showed that decorin can inhibit the high glucose-induced oxidative stress in lens epithelial cells in vitro. The administration of decorin decrease the ROS production, increase the ratio of GSH/GSSG and increase the expression ratio of bcl-2 to bax in cells treated with high glucose, which suggests decorin may inhibit the development of diabetic cataracts. Moreover, decorin can inhibit p22phox induction and p38 MAPK activation of HLEB3 cells under high glucose condition. However, we only investigated the inhibitory influence of decorin on p22phox-p38 MAPK pathway. We did not explore the other possible pathways and the downstream mechanism following the inhibition of p38 MAPK was not explored also. For financial reasons, we only used human len epithelium cell line HLEB3 to study. Further research is needed to illuminate the other mechanisms of decorin on the oxidative stress of HLE cells in high glucose condition with primary cells.

In a word, our study proved that decorin can reduce the increment of oxidative stress injury and the apoptotic ratio of HLE cells significantly under high glucose condition in vitro. This effect was mainly regulated suppressing the activation of p22phox-p38 MAPK pathway induced by high glucose. These findings suggested that decorin may have beneficial influence on the development of diabetic cataract.

## Author contribution statements

Fengyan Zhang and Shanshan Du conceived the study. Jingzhi Shao and Dandan Xie performed the experiments. Shanshan Du and Jingzhi Shao analyzed the data. Shanshan Du and Jingzhi Shao drafted the manuscript. All authors read and approved the final version of manuscript.

## Acknowledgments

This work was funded by the scientific and technological project of Henan Province (No.192102310343), medical scientific and technological project of Henan Province (No.2018020065), National Natural Science Foundation of China (81970785) and the First Affiliated Hospital of Zhengzhou University.

The authors thank Professor Fuguang Li (the laboratory of Department of Immunology, school of Basic Medical, Zhengzhou University) for providing the standardized experimental platform; Lili Jiang (school of Basic Medical, Zhengzhou University) for assistant with flow cytometry analysis.

## Ethics statement

Patients involved here have not subjected to any specific clinical treatments and have all signed informed cosent. This study was approved by the Ethics Committee of the First Affiliated Hospital of Zhengzhou University.

